# SAMHD1 promotes oncogene-induced replication stress

**DOI:** 10.1101/2020.07.29.226282

**Authors:** Si Min Zhang, Jose M Calderón-Montaño, Sean G Rudd

**Author notes:** Department of Pharmacology, Faculty of Pharmacy, University of Seville, Spain (Jose M Calderón-Montaño).

## Abstract

Oncogenes induce DNA replication stress in cancer cells. Although this was established more than a decade ago, we are still unravelling the molecular underpinnings of this phenomenon, which will be critical if we are to exploit this knowledge to improve cancer treatment. A key mediator of oncogene-induced replication stress is the availability of DNA precursors, which will limit ongoing DNA synthesis by cellular replicases. In this study, we identify a potential role for nucleotide catabolism in promoting replication stress induced by oncogenes. Specifically, we establish that the dNTPase SAMHD1 slows DNA replication fork speeds in human fibroblasts harbouring an oncogenic RAS allele, elevating levels of endogenous DNA damage, and ultimately limiting cell proliferation. We then show that oncogenic RAS-driven tumours express reduced SAMHD1 levels, suggesting they have overcome this tumour suppressor barrier, and that this correlates with worse overall survival for these patients.

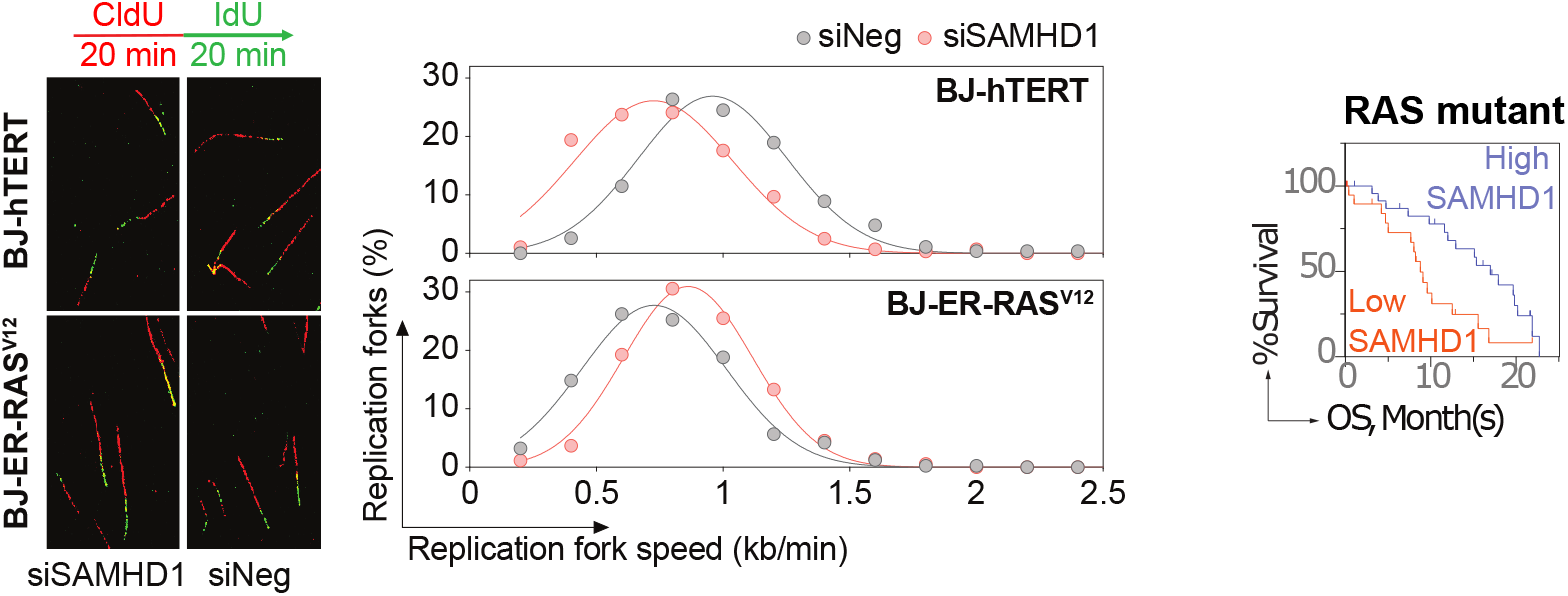

## Introduction

One-third of human cancers contain mutations in the RAS gene family, placing the resultant oncogenes amongst the most frequent drivers of tumour development (1). A central feature of these tumours is the presence of DNA replication stress (RS) (2), which is when DNA synthesis slows or stalls but DNA unwinding continues, resulting in exposed stretches of single-stranded DNA. If left unattended, this can lead to the formation of DNA breaks as well as aberrant mitoses, in turn promoting genome instability linked to disease aggressiveness and therapy resistance. Yet, it can also be a mechanism of tumour suppression, as it slows cell proliferation and activates anticancer barriers such as apoptosis and senescence (3). Oncogene-induced RS is thus a double-edged sword, and a complete understanding of the molecular mechanisms underpinning this process, together with the factors involved, will be central to the identification of potential biomarkers and therapeutic targets that can be utilised to improve the treatment of these cancers.

There are several mechanisms by which oncogenes such as RAS induce RS during early cancer development (2), one of which is by limiting the availability of deoxynucleoside triphosphates (dNTPs) (4). For instance, elevated origin firing, and the resulting increase in DNA synthesis, will exhaust dNTP pools at a faster rate. Oncogenic RAS signalling also downregulates key dNTP biosynthetic enzymes such as ribonucleotide reductase and thymidylate synthase (5,6), whilst the remaining purine dNTPs can become oxidised and subsequently sanitised from the dNTP pool (7). All of this is underscored by oncogene-induced RS in cultured cells being rescued by supplementation of the culture medium with nucleosides (5,6,8,9) or folate (10), a micronutrient required for nucleotide biosynthesis. Following induction of RS by oncogenes, ATR-Chk1 signalling will be activated, which is largely responsible for stabilising and restarting stalled forks (2), and can accordingly protect cancer cells from oncogene-induced RS in a concentration-dependent manner (11).

In the present study, we hypothesised the involvement of SAM and HD-domain containing protein-1 (SAMHD1) in oncogene-induced RS for two reasons. Firstly, SAMHD1 is a dNTP triphosphohydrolase positioned as a central regulator of dNTP pools (12,13), and thus this catabolic activity could potentially limit available dNTPs in cancer cells harbouring oncogenes such as RAS. Secondly, SAMHD1 has a non-catalytic role in ATR-Chk1 activation and subsequent replication fork restart (14), and this role could potentially protect tumour cells from oncogene-induced RS.

## Results and Discussion

To investigate the potential role of SAMHD1 in oncogene-induced RS, we began by utilising a well-described defined genetic system (15). In this system, normal human BJ fibroblasts have been transformed in a stepwise manner, first using telomerase catalytic subunit (hTERT), to generate the immortalized but non-tumorigenic BJ-hTERT cell line, and subsequently with Simian Virus 40 (SV40) Early Region (ER) (encompassing small and large T antigen (16,17)) and an allele of oncogenic RAS (*HRAS*^V12^), ultimately creating BJ-ER-RAS^V12^ cells, which have lethal metastatic potential (15) (**Fig. 1A**). Expression of RAS^V12^ in cultured cells induces cell hyperproliferation followed by RS and RS-induced senescence (3,5,6,9), however the presence of SV40 ER prevents senescence onset and thereby phenotypically locks this cell model in a hyperproliferative state (15–17). We first confirmed an elevated proportion of replicating cells following oncogene expression by comparing cell cycle profiles between BJ-hTERT and BJ-ER-RAS^V12^ cells (**Fig. 1B**), and furthermore, confirmed the presence of replication stress by measuring the speeds of individual replication forks and endogenous DNA damage present in these cell lines (discussed in detail below). Intriguingly, when assessing SAMHD1 abundance in these cell lines, we observed that BJ-ER-RAS^V12^ cells exhibited elevated levels of SAMHD1 transcript (**Fig. 1C**) and protein (**Fig. 1D)**compared to BJ-hTERT cells. Furthermore, compared to >40 cancer cell lines, BJ-ER-RAS^V12^ cells exhibited the highest SAMHD1 transcript abundance, followed by BJ-hTERT cells transformed with SV40 ER (BJ-hTERT-ER) (**Supplementary Fig.1**), supporting a potential role for SAMHD1 in the context of oncogene expression

**Figure 1.**
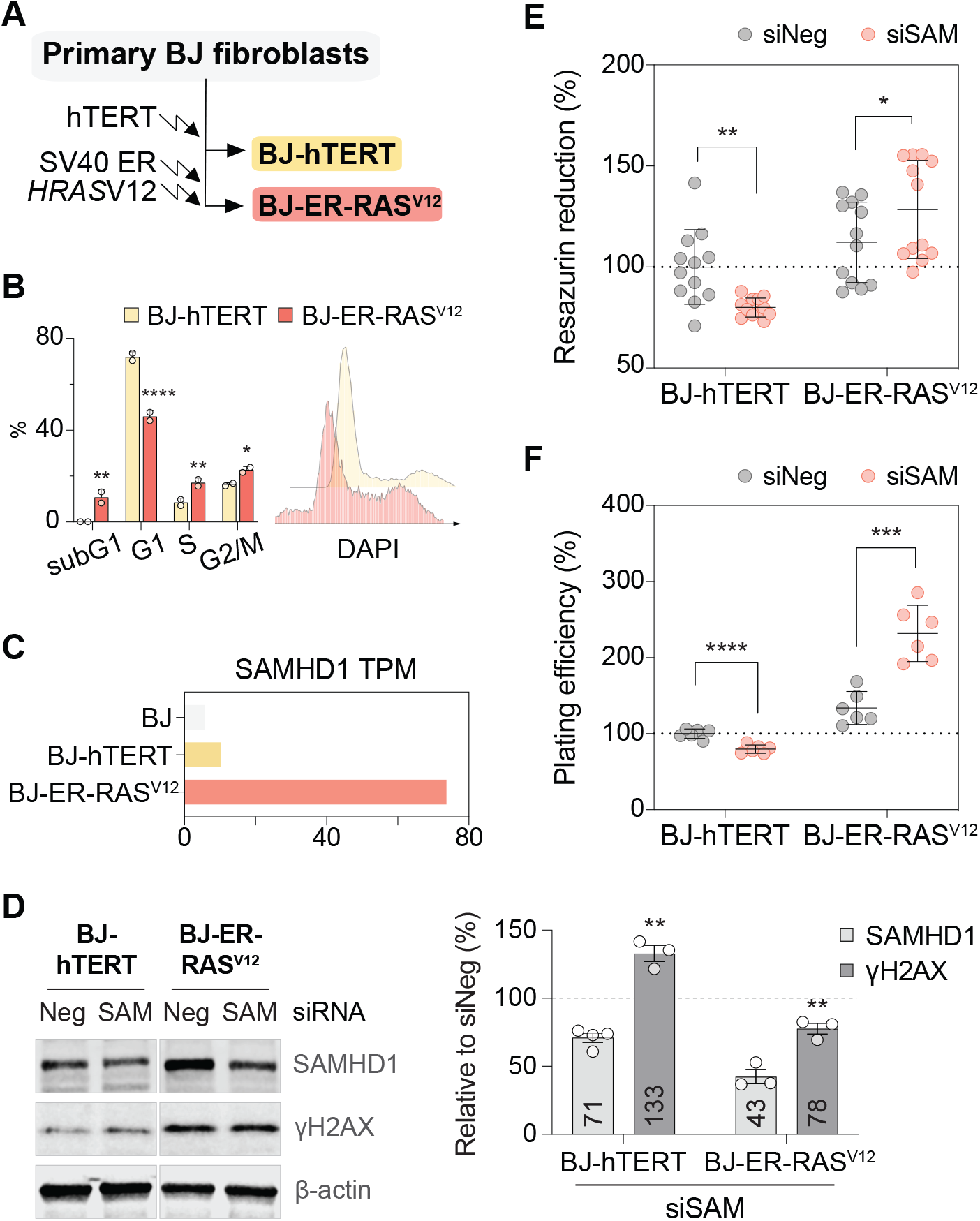
SAMHD1 limits the proliferation of cells harbouring RAS^V12^. **A.** Schematic detailing generation of BJ-hTERT and BJ-ER-RAS^V12^ cells. **B.** Cell cycle profiles generated by high-content imaging of nuclear DNA content. Left panel: cell cycle profile analysis of cells stained with DAPI followed by high-content imaging of nuclear DNA content. Data summarized from 2 independent experiments, each with >10,000 nuclei analysed per cell line; BJ-hTERT Vs. BJ-ER-RAS^V12^, * *p ≤* 0.05, ** *p* ≤ 0.01, **** *p* ≤ 0.0001 (Two-way ANOVA, Fisher’s LSD test). Right panel: histograms from a representative experiment shown. **C.** SAMHD1 transcripts per million (TPM). **D.** Western blot analysis of cell lysates 72 hours post-transfection with SAMHD1-targeting (siSAM) or control (siNeg) siRNA. Left panel: Representative Western blot image. Right panel: densitometry analysis of repeat experiments (BJ-hTERT, n=4; BJ-ER-RAS^V12^, n=3 total experiments analysed). Protein levels were normalised to siNeg and indicated in the graph; siNeg Vs. siSAM, ** *p* ≤ 0.01 (Student’s t-test, two-tailed, unpaired). **E,F.** Proliferation analysis of siRNA-transfected cells as in (D) by resazurin assay (E) or colony formation assay (F), re-seeded 48 hours post-transfection and assessed at 4 or 10 days post-seeding, respectively. Values relative to BJ-hTERT siNeg (dashed line) plotted. Individual replicates of 2 independent experiments shown, mean ± s.d. indicated; siNeg Vs. siSAM, * *p* ≤ 0.05, ** *p* ≤ 0.01 (Student’s t-test, unpaired).

We reasoned that elevated SAMHD1 levels could either contribute to oncogene-induced RS by limiting dNTP supply or counteract oncogene-induced RS by promoting restart of stalled replication forks. To investigate these putative roles, we utilised a pool of four SAMHD1-specific siRNAs and transfected the BJ-hTERT and BJ-ER-RAS^V12^ cells, achieving a partial knockdown of SAMHD1 protein (**Fig. 1D**). Consistent with RS induction by RAS^V12^, an elevated level of DNA damage marker phosphorylated histone variant γH2AX was present in the BJ-ER-RAS^V12^ cells compared to BJ-hTERT cells (**Fig. 1D**). Interestingly, whilst siRNA-mediated reduction of SAMHD1 level in the BJ-hTERT cells resulted in elevated γH2AX signal, in BJ-ER-RAS^V12^ cells, a mild reduction was observed (**Fig. 1D**), suggesting a role for SAMHD1 in promoting RAS^V12^-induced DNA damage in these cells.

Given oncogene-induced RS can limit cell proliferation, we next monitored this phenotype using a short-term metabolic cell viability assay and a long-term colony formation assay. Depletion of SAMHD1 in BJ-hTERT cells led to a small, but significant, reduction in the proportion of cells in both assays (**Fig. 1E, F**), concordant with the increase in γH2AX signal induction (**Fig. 1D**). In contrast, SAMHD1 depletion in BJ-ER-RAS^V12^ cells resulted in a significant increase in the proportion of cells (**Fig 1E, F**). This was particularly striking in the colony formation assay in which a doubling of plating efficiency following SAMHD1 depletion was observed (**Fig. 1F**). Given BJ-ER-RAS^V12^ cells express SV40 ER, as a control experiment, we also depleted SAMHD1 in BJ-hTERT cells transformed only with SV40 ER (BJ-hTERT-ER cells) and monitored cell growth. However, no difference in cell viability was observed following SAMHD1 depletion (**Supplementary Fig. 2**) despite a report indicating a role for SAMHD1 in telomere maintenance following SV40 large T-transformation (18).

To directly examine RS, we next measured the speeds of individual replication forks in the siRNA-treated BJ-hTERT and BJ-ER-RAS^V12^ cells using the DNA fibre assay (**Fig. 2A**). BJ-ER-RAS^V12^ cells had slower fork speeds than BJ-hTERT cells (**Fig. 2B, C**), consistent with RAS^V12^-induced RS. Notably, whilst depletion of SAMHD1 in BJ-hTERT cells led to a reduction in fork speeds, the opposite response was observed in BJ-ER-RAS^V12^ cells, with a significant increase in fork speeds being apparent (**Fig. 2B, C**). Taken together, these data demonstrate that SAMHD1 limits the proliferation of cells harbouring RAS^V12^ by slowing DNA replication fork speeds.

**Figure 2.**
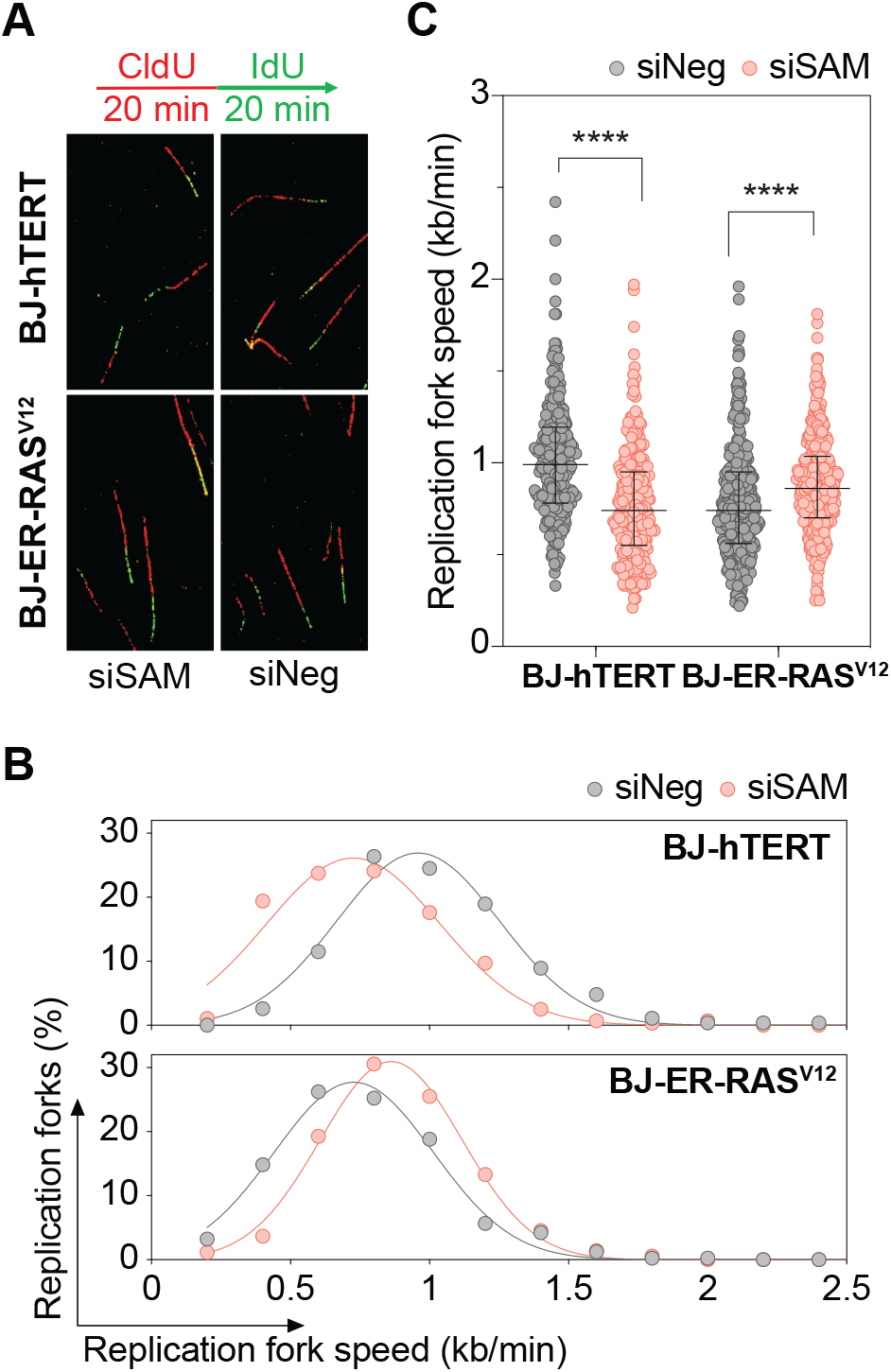
SAMHD1 slows replication fork speeds in cells harbouring RAS^V12^. **A.** Representative images of spread DNA fibres from BJ-hTERT and BJ-ER-RAS^V12^ cells 72 hours post-transfection with SAMHD1-targeting (siSAM) or control (siNeg) siRNA. Assay schematic shown above. **B.** Histogram of DNA replication fork speeds. Values derived from 2 independent experiments. Total DNA fibres analysed: BJ-hTERT siNeg, 269; siSAM, 278; BJ-ER-RAS^V12^ siNeg, 404; siSAM, 353. **C.** Individual DNA replication fork speeds with median ± interquartile range shown from 2 independent experiments; siNeg Vs. siSAM, *** *p* ≤ 0.001, **** *p* ≤ 0.0001 (two-tailed Mann Whitney test).

RAS mutations have long been established as the drivers of multiple malignancies, and our data suggests that SAMHD1 loss-of-function and/or downregulation, which occurs in a number of different malignancies (13), could confer a particular proliferative advantage to oncogenic Ras-driven tumour clones, contributing to poorer patient survival. Thus, utilising data from The Cancer Genome Atlas (TCGA), we next interrogated SAMHD1 expression in malignancies most commonly driven by oncogenic RAS mutations (**Supplementary Fig. 3**). Presence of an oncogenic KRAS allele in pancreatic (PAAD) (mutation rate, 59.2%), colorectal (COAD) (36.2%), and lung adenocarcinoma (LUAD) (26.7%), corresponded to significantly lower levels of SAMHD1 mRNA when compared to their KRAS wild-type (WT) counterparts (**Fig. 3A**). A similar trend was observed with NRAS-driven tumours (**Fig. 3B**), but to a lesser degree for HRAS (**Fig. 3C**), which typically have lower mutation frequencies of below 27.9% and 10.1%, respectively. Critically, those patients with oncogenic KRAS-driven tumours and low SAMHD1 expression had a worse overall survival (OS) when compared to those with high SAMHD1 expression, in stark contrast to their WT KRAS counterparts which had similar OS regardless of SAMHD1 levels (**Fig. 4A**, **B**). This trend was particularly striking in KRAS-driven PAAD (**Fig. 4B**). Overall, these data suggest that SAMHD1 is a tumourigenic barrier in oncogenic RAS-driven malignancies, and accordingly, suggest overcoming this barrier may lead to a proliferation advantage for cancer cells and ultimately worse OS.

**Figure 3.**
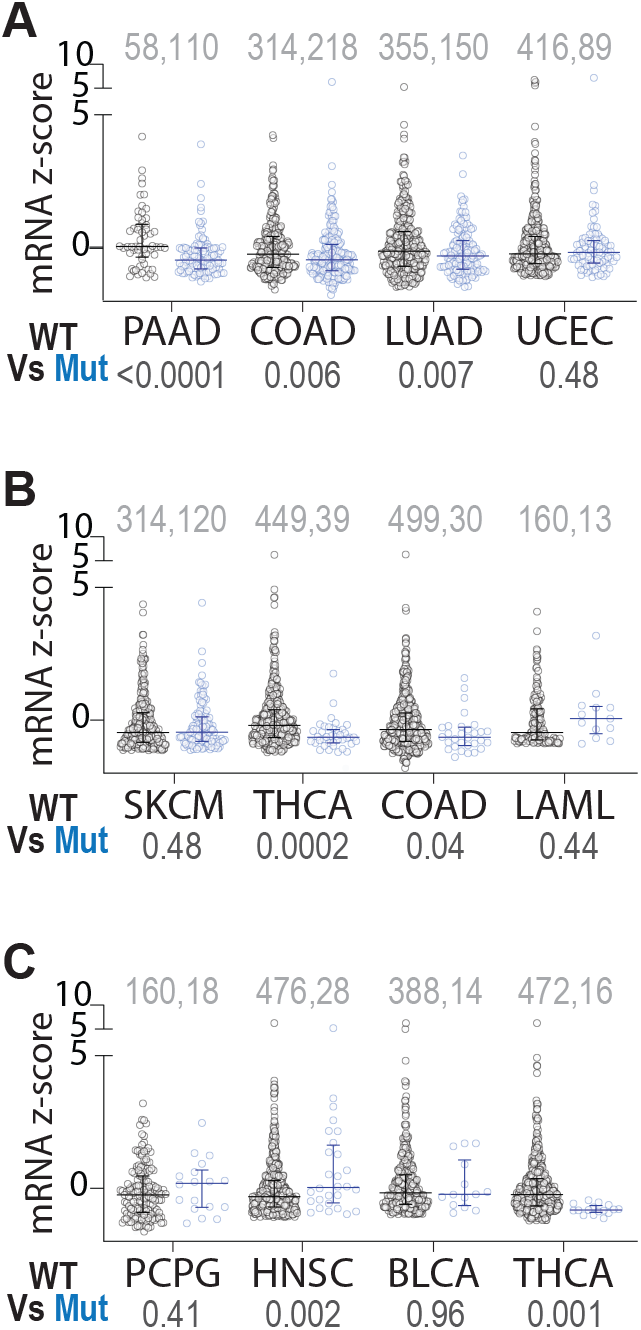
Oncogenic RAS-driven tumours downregulate SAMHD1. SAMHD1 mRNA levels (z-scores) in TCGA tumour samples containing wildtype (WT) or oncogenic (Mut) KRAS (**A**), NRAS (**B**) or HRAS (**C**). Individual sample value with median ± interquartile range shown, n and *p* values (calculated using unpaired t test) indicated above and below, respectively.

**Figure 4.**
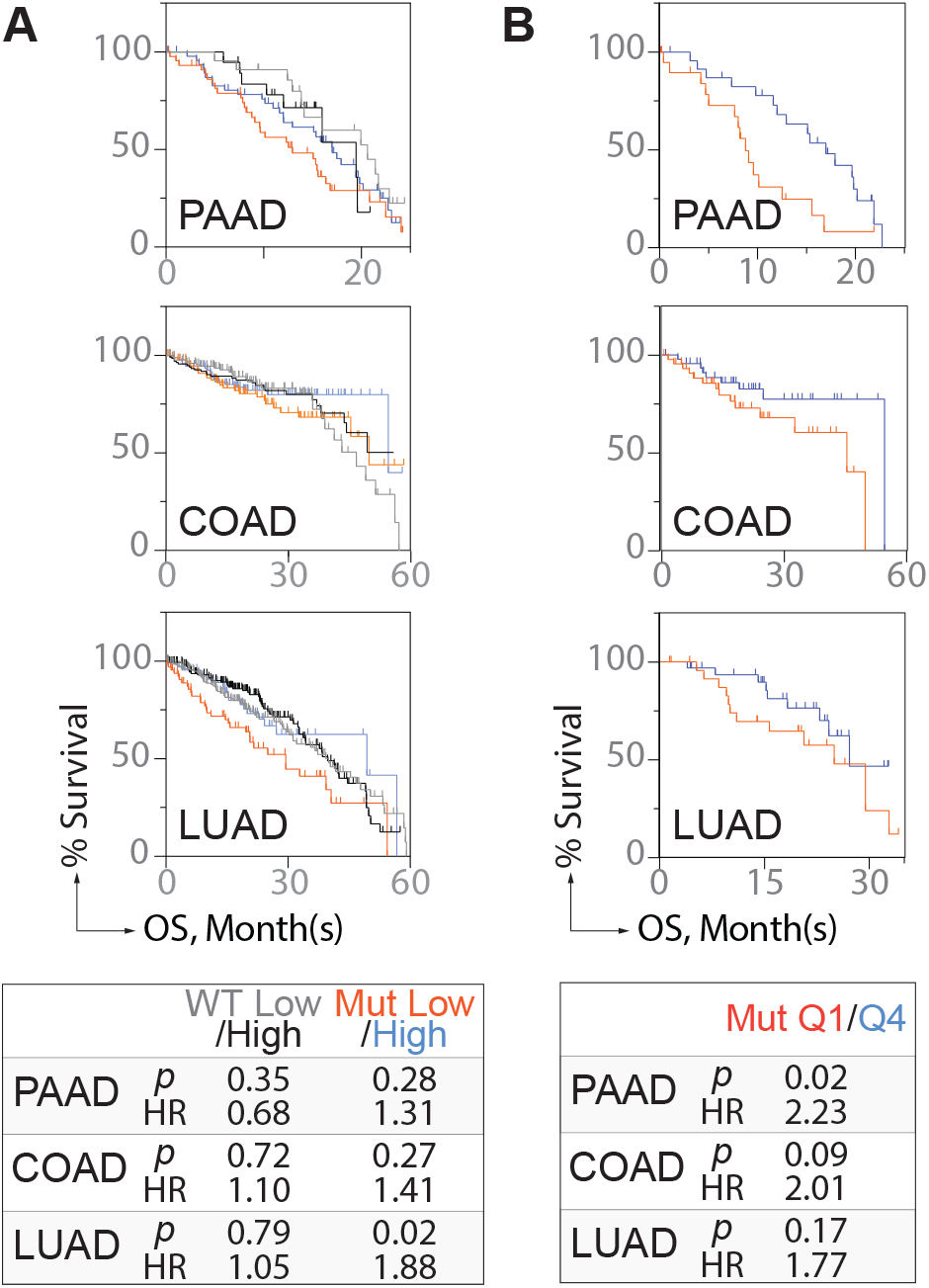
Oncogenic-RAS driven malignancies with low SAMHD1 expression are associated with worse overall survival. A. Kaplan–Meier overall survival (OS) analysis of TCGA pancreatic (PAAD), colon (COAD), and lung adenocarcinoma (LUAD) patients harbouring WT or oncogenic (Mut) KRAS, after dichotomization for SAMHD1 mRNA levels (z-score) for below (PAAD, n=22(WT), 45(Mut); COAD, n=131(WT), 94(Mut); LUAD, n=157(WT), 67(Mut)) and above (PAAD, n=19(WT), 49(Mut); COAD, n=141(WT), 96(Mut); LUAD, n=152(WT), 68(Mut)) median. SAMHD1 mRNA median values, WT/Mut: PAAD, 0.0374/-0.5292; COAD, −0.2638/-0.4725; LUAD, −0.1579/-0.3399. B. Kaplan–Meier OS analysis of cohorts in (A) with mutant KRAS-expressing tumours, with bottom (PAAD, n=20; COAD, n=45; LUAD, n=30) and top 25% (PAAD, n=25; COAD, n=48; LUAD, n=35) SAMHD1 mRNA levels. HR, hazard ratio.

In this study we set out to test the hypothesis that SAMHD1 is involved in oncogene-induced RS. We speculated this could be due to the catabolic role of SAMHD1 in dNTP homeostasis and/or the role in replication fork restart; our data indicates the former could play a more prominent role in this context, although further work is required to establish the precise mechanism. There is a well-established link between reduced dNTPs and oncogene-induced RS in early cancer development (2,4). Oncogenic signalling will elevate origin firing, consuming available dNTPs at a faster rate, whilst downregulating nucleotide biosynthetic enzymes (5,6) and inducing oxidation of the dNTP pool (7), all of which will reduce available dNTPs needed for on-going DNA synthesis. Accordingly, oncogene-induced RS in cultured cells can be reverted by nucleoside supplementation (5,6,8,9). The data presented here indicates that a reduction of dNTP catabolism via downregulation of the dNTPase SAMHD1, a well-documented consequence of which is expansion of the dNTP pool (12,13), can rescue oncogene-induced RS. The data presented here also indicates that this tumour suppressor role of SAMHD1 is clearly dictated by the oncogenic re-wiring present in cancer cells, offering an explanation for the differing proliferative phenotypes reported from loss of SAMHD1 and/or dNTP pool expansion (13).

To achieve sustainable long-term growth, oncogene-driven malignancies need to maintain sufficient dNTP pools, and this can be achieved through metabolic rewiring. This has been demonstrated in the case of oncogenic BRAF (19) and KRAS-driven PAAD (20), which upregulate de novo dNTP synthesis via the pentose phosphate pathway. Our analysis of clinical data suggests downregulation of dNTP catabolism via SAMHD1 could be an important component of this rewiring. Critically, under these circumstances, this corresponds to worse patient survival. Altogether, this study establishes SAMHD1 as a tumourogenic barrier in these RAS-driven malignancies through promoting RS, and opens further avenues of investigation regarding the interplay of dNTP catabolism and oncogenesis.

## Materials and Methods

### Cell culture

Genetically modified normal human BJ fibroblast lines BJ-hTERT (BJ cells immortalised with hTERT), BJ-hTERT-ER (SV40 Early Region transformed BJ-hTERT cells), and BJ-ER-RAS^V12^ (HRAS^V12^-transformed BJ-hTERT-ER cells) were provided by William C. Hahn (Dana Faber Cancer Institute) and their generation has been reported previously (15). Of note, whilst the original publication referred to these cell lines being transformed with SV40 Large-T antigen (15), a later study clarified that they were instead constructed with the entire SV40 ER, encompassing both small T and large T (16). Cells were cultured in DMEM supplemented with 10% FBS and penicillin-streptomycin at 37°C with 5% CO2 in a humidified incubator. All cell culture reagents were purchased from Gibco/Thermo Fisher. Cells were typically maintained in culture for 1-2 months before thawing of a new batch and mycoplasma screening was performed routinely using the MycoAlert Mycoplasma Detection Kit (Lonza).

### RNA interference

Transfections were performed using INTERFERin (Polyplus Transfection) following manufacturer’s instructions. SAMHD1-targeting siRNA pool (siGENOME SMARTpool; M-013950-00-0005, Dharmacon) or control siRNA (AllStars Negative Control, Qiagen) were transfected at 10 nM final concentration.

### Western blot analysis

Cells were scraped in lysis buffer (50 mM Tris-HCl pH 8, 150 mM NaCl, 1 mM EDTA, 1% Triton X-100, 0.1% SDS) supplemented with cOmplete EDTA-free protease inhibitor cocktail (Roche) and Halt phosphatase inhibitor cocktail (Thermo Fisher). Samples were incubated on ice for 30 mins to 1 hour with occasional vortexing before centrifugation to pellet insoluble material. Protein concentration of the remaining soluble fraction was determined using the Pierce BCA Protein Assay Kit (Thermo Fisher) and samples with equal total protein quantity were prepared with Laemmli Sample Buffer (Bio-Rad) before denaturation at 95 °C for 5 min. SDS–PAGE was performed using a Bio-Rad setup with Criterion TGX 4–20% gels (Bio-Rad) and proteins were transferred to a nitrocellulose membrane using Trans-Blot Turbo Transfer System (Bio-Rad), all according to the manufacturer’s instructions. Membranes were blocked in Odyssey Blocking Buffer (Li-Cor) and probed with primary and then species-appropriate IRDye-conjugated secondary antibodies (Li-Cor) before visualization on an Odyssey Fc Imaging System (Li-Cor). Primary antibodies used in this study were anti-SAMHD1 (Abcam, ab128107; 1:1000), anti-γH2AX (Millipore, 05-636, 1:2000) and anti-β-actin (Abcam, ab6276; 1:5000). Densitometry analysis on acquired Western Blot images was performed using Image Studio Lite (Ver. 5.2.; Li-Cor), with signals subsequently normalised to β-actin signals and relative to that of siNeg-treated cells.

### High content microscopy

Cells were seeded (10,000 per well) in black clear-bottomed 96-well plates (BD Falcon) and allowed to grow for 24 hours. Cells were then fixed with 4% paraformaldehyde in PBS (Santa Cruz) supplemented with 0.5% Triton X-100 for 15 minutes. Wells were washed with PBS before staining cell nuclei with 1 μg/ml DAPI for 10 minutes. Wells were washed again with PBS and imaged on an ImageXpress high-throughput microscope (Molecular Devices) with a 20x objective. Images were analysed (nuclei counting and DAPI intensity measurements) with CellProfiler (21) (Broad Institute) and data handled in Excel (Microsoft) and plotted in Prism 8 (GraphPad).

### Resazurin assay

Cells (48 hrs post siRNA transfection) were seeded into 96-well plates with 1500 cells per well in 100 μl medium and incubated for 4 days. Following incubation, resazurin (Sigma Aldrich, cat no. R7017) stock solution (1 mg/ml in PBS) was diluted in fresh medium and 100 μl added to wells to a final concentration of 0.1 mg/ml, and incubated for 2 hours. Resazurin reduction to resorufin, an indicator of cell viability, was subsequently measured at 530/590 nm (ex/em) using a Hidex Sense Microplate Reader. Fluorescent intensity for each well was normalised to the average of BJ-hTERT cell control wells on the same plate to give relative cell viability values.

### Colony formation assay

Cells (48 hrs post siRNA transfection) were seeded onto 10 cm dishes and incubated for 10 days. For BJ-hTERT, 1000 cells were seeded per dish, whilst BJ-hTERT-ER and BJ-ER-RAS^V12^, 500 cells were seeded per dish. Following incubation, colonies were fixed and stained with 4% methylene blue in methanol. Colonies were counted manually, and plating efficiency ((no. of colonies formed/no. of cells plated) x 100) calculated and normalized to the average of BJ-hTERT cell control wells.

### DNA fibre assay

Cells (48 hrs post siRNA transfection) were seeded into 6-well plates to reach 70% confluency the following day, and then pulse-labelled for 20 minutes with 5-Chloro-deoxyuridine (CldU, Sigma) and subsequently 20 minutes with 250 μM 5-Iodo-deoxyuridine (IdU, Sigma). Cells were collected and re-suspended in ice-cold PBS before lysing and incubating a drop of cell suspension together with spreading buffer (200 mM Tris-HCl pH 7.4, 50 mM EDTA, 0.5% SDS) on SuperFrost objective glass slides (Thermo Fisher). Slides were tilted to allow spreading of the DNA fibres. For immunodetection of CldU/IdU, acid treated fibres (2.5 M HCl for 1 hour) were stained with monoclonal rat anti-BrdU (1:500, Clone BU1/75 (ICR1), AbD Serotec) and monoclonal mouse anti-BrdU (1:500, Clone B44, 347580, BD Biosciences) respectively for 1 hour at 37 °C. Subsequently, slides were incubated with Alexa 555-conjugated anti-rat and Alexa 488-conjugated anti-mouse secondary antibodies (Molecular Probes). DNA fibres were imaged in a LSM780 confocal microscope using a 64x oil objective and lengths of CldU and IdU tracks were measured using ImageJ software (http://fiji.sc/Fiji).

### Expression and survival analyses

Cell line RNA-sequencing data was obtained from The Human Protein Atlas project (https://www.proteinatlas.org) (22) and plotted in Prism 8 (GraphPad). Tumour mRNA expression and patient survival data was obtained via cBioPortal (23,24) from TCGA PanCancer Atlas cohorts (25), and plotted in Prism 8 (GraphPad).

### Statistical testing

Statistical testing was performed in Prism 8 (GraphPad), with the parameters for each main figure listed below, and those for the supplementary figures included in their respective legends.

For comparison of cell cycle profiles between BJ-hTERT and BJ-ER-RAS^V12^ cells in Fig. 1a, statistical significance was determined using two-way ANOVA, Fisher’s LSD test: subG1, *p* = 0.0013, t = 3.654, DF = 12; G1, *p* < 0.0001, t = 10.26, DF = 12; S, *p* = 0.0055, t = 3.379, DF = 12; G2/M, *p* = 0.0270, t = 2,517, DF = 12.

For comparison of γH2AX level upon siSAM-mediated SAMHD1 knockdown in Fig. 1d, statistical significance was determined using a two-tailed unpaired Student’s t-test: siNeg vs siSAM in BJ-hTERT cells, *p* = 0.005126, t ratio = 5.559, df = 4; in BJ-ER-RAS^V12^ cells, *p* = 0.005417, t ratio = 5.475, df = 4.

For comparison of cell proliferation upon siSAM-mediated SAMHD1 knockdown in Fig. 1e, statistical significance was determined using a two-tailed unpaired Student’s t-test: siNeg vs siSAM in BJ-hTERT cells, *p* = 0.003, t ratio = 3.042, df = 66; in BJ-ER-RAS^V12^ cells, *p* = 0.016, t ratio = 2.477, df = 66.

For comparison of colony formation upon siSAM-mediated SAMHD1 knockdown in Fig. 1f, statistical significance was determined using a two-tailed unpaired Student’s t-test: siNeg vs siSAM in BJ-hTERT cells, *p* = 0.0001, t ratio = 6.105, df = 10; in BJ-ER-RAS^V12^ cells, *p* = 0.0002, t ratio = 5.613, df = 10.

For comparison of replication fork speeds upon siSAM-mediated SAMHD1 knockdown in Fig. 2c, statistical significance was determined using a two-tailed Mann Whitney test: siNeg vs siSAM in BJ-hTERT cells, *p* < 0.0001, U = 20885; in BJ-ER-RAS^V12^ cells, *p* < 0.0001, U = 53025.

For comparison of SAMHD1 mRNA levels in tumours containing WT RAS vs Mut RAS in Fig. 3, statistical significance was determined using one-way ANOVA Fisher’s LSD test: Fig. 3a, WT KRAS vs Mut KRAS (PAAD, *p* < 0.0001, t = 4.698, DF = 377; COAD, *p* = 0.0058, t = 2.768, DF = 880; LUAD, *p* = 0.0067, t = 2.717, DF = 801; UCEC, *p* = 0.4767, t = 0.7120, DF = 688); Fig. 3b, WT NRAS vs Mut NRAS (SKCM, *p* = 0.4831, t = 0.7017, DF = 669; THCA, *p* = 0.0002, t = 3.776, DF = 561; COAD, *p* = 0.0372, t = 2.088, DF = 581; LAML, *p* = 0.4368, t = 0.7793, DF = 190); Fig. 3c, WT HRAS vs Mut HRAS (PCPG, *p* = 0.4068, t = 0.8313, DF = 207; HNSC, *p* = 0.0017, t = 3.147, DF = 554; BLCA, *p* = 0.9585, t = 0.05212, DF = 427; THCA, *p* = 0.001, t = 3.317, DF = 515).

For comparison of OS of patients with tumours expressing low vs high levels of SAMHD1 in Fig. 4a, Kaplan-Meier survival analyses using Mantel-Cox log-rank test was used: in PAAD patients (with WT KRAS background, n = 22 and 19, respectively, *p* = 0.3513, *χ2* = 0.8687, df =1, HR = 0.6753; with Mut KRAS background, n = 45 and 49, respectively, *p* = 0.2779, *χ2* = 1.177, df =1, HR = 1.312); in COAD patients (with WT KRAS background, n = 131 and 141, respectively, *p* = 0.7163, *χ2* = 0.1320, df =1, HR = 1.1; with Mut KRAS background, n = 94 and 96, respectively, *p* = 0.2735, *χ2* = 1.199, df =1, HR = 1.412); in LUAD patients (with WT KRAS background, n = 157 and 152, respectively, *p* = 0.7890, *χ2* = 0.07163, df =1, HR = 1.050; with Mut KRAS background, n = 67 and 68, respectively, *p* = 0.0236, *χ2* = 5.123, df =1, HR = 1.876).

For comparison of OS of patients with Mut KRAS tumours expressing low (Q1) vs high (Q4) levels of SAMHD1 in Fig. 4b, Kaplan-Meier survival analyses using Mantel-Cox log-rank test was used: in PAAD patients, n = 20 and 25, respectively, *p* = 0.0159, *χ^2^* = 5.812, df =1, HR = 2.227; in COAD patients, n = 45 and 48, respectively, *p* = 0.0896, *χ^2^* = 2.882, df =1, HR = 2.006; in LUAD patients, n = 30 and 35, respectively, *p* = 0.1697, *χ^2^* = 1.886, df =1, HR = 1.773.

## Data availability

Materials, protocols, and source data used in this study are available from the corresponding author upon request.

## Acknowledgements

We thank Thomas Helleday and Nicholas Valerie for comments on the data and manuscript. Results shown here are in part based upon data generated by the TCGA Research Network (https://www.cancer.gov/tcga). This work was supported by the Felix Mindus contribution to Leukemia Research (2019-02004 to S.M.Z.), the Swedish Research Council (2018-02114 to S.G.R), the Swedish Cancer Society (19-0056-JIA to S.G.R), and the Swedish Childhood Cancer Foundation (PR2019-0014 to S.G.R).

## Author contributions

S.G.R. and S.M.Z. conceptualised the study. S.M.Z., J-M.C.M. and S.G.R. designed, performed, and analysed experiments. S.M.Z. and S.G.R. prepared the figures and wrote the paper. S.G.R. supervised the study.

## Conflict of interest

The authors have no competing financial interests in relation to the work described.

## Supplementary Figures

**Supplementary Figure 1.**
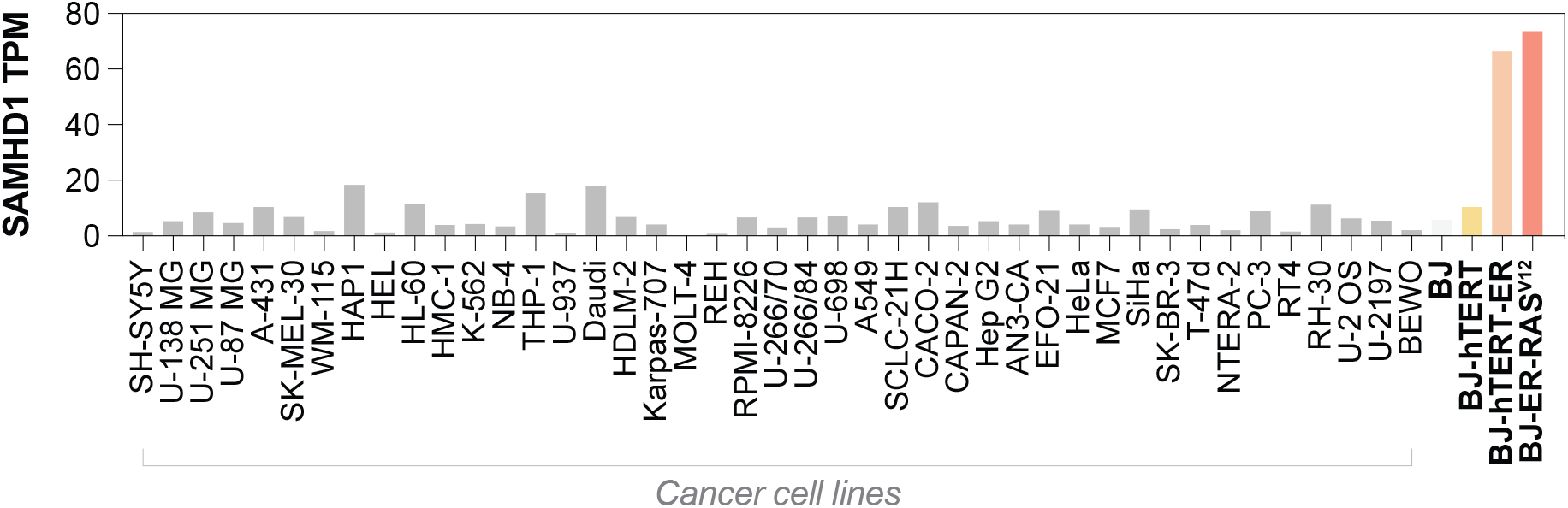
SAMHD1 transcripts per million (TPM) in BJ-hTERT, BJhTERT-ER, and BJ-ER-RAS^V12^ cells, together with a panel of 43 cancer cell lines. Data extracted from The Human Protein Atlas project (https://www.proteinatlas.org) (22).

**Supplementary Figure 2.**
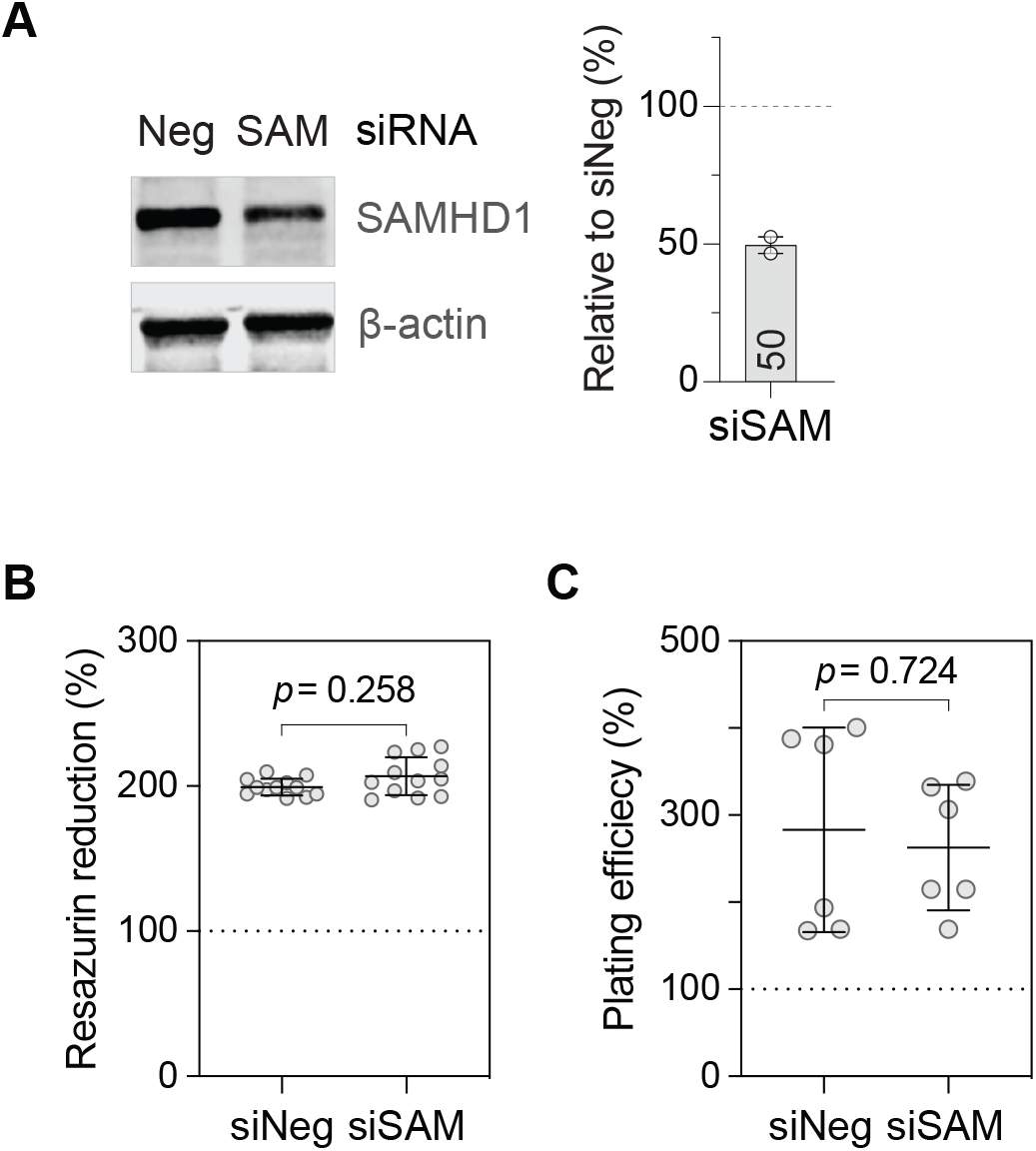
SAMHD1 depletion does not affect the proliferation of BJ-hTERT-ER cells. **A.** Western blot analysis of cell lysates 72 hours post-transfection with SAMHD1-targeting (siSAM) or control (siNeg) siRNA. Left panel: Representative Western blot images. Right panel: densitometry analysis of repeat experiments (n=2 total experiments analysed). Protein levels were normalised to siNeg and indicated in the graph. **B,C.** Proliferation analysis of siRNA-transfected cells as in (A) by resazurin assay (B) or colony formation assay (C), re-seeded 48 hours post-transfection and assessed at 4 or 10 days post-seeding, respectively. Values relative to BJ-hTERT siNeg (dashed line) plotted. Individual replicates of 2 independent experiments shown, mean ± s.d. and *p* values (Student’s t-test, unpaired) indicated. Statistical testing information: for (B), t ratio = 1.141, df = 66; for (C), t ratio = 0.362, df = 10.

**Supplementary Figure 3.**
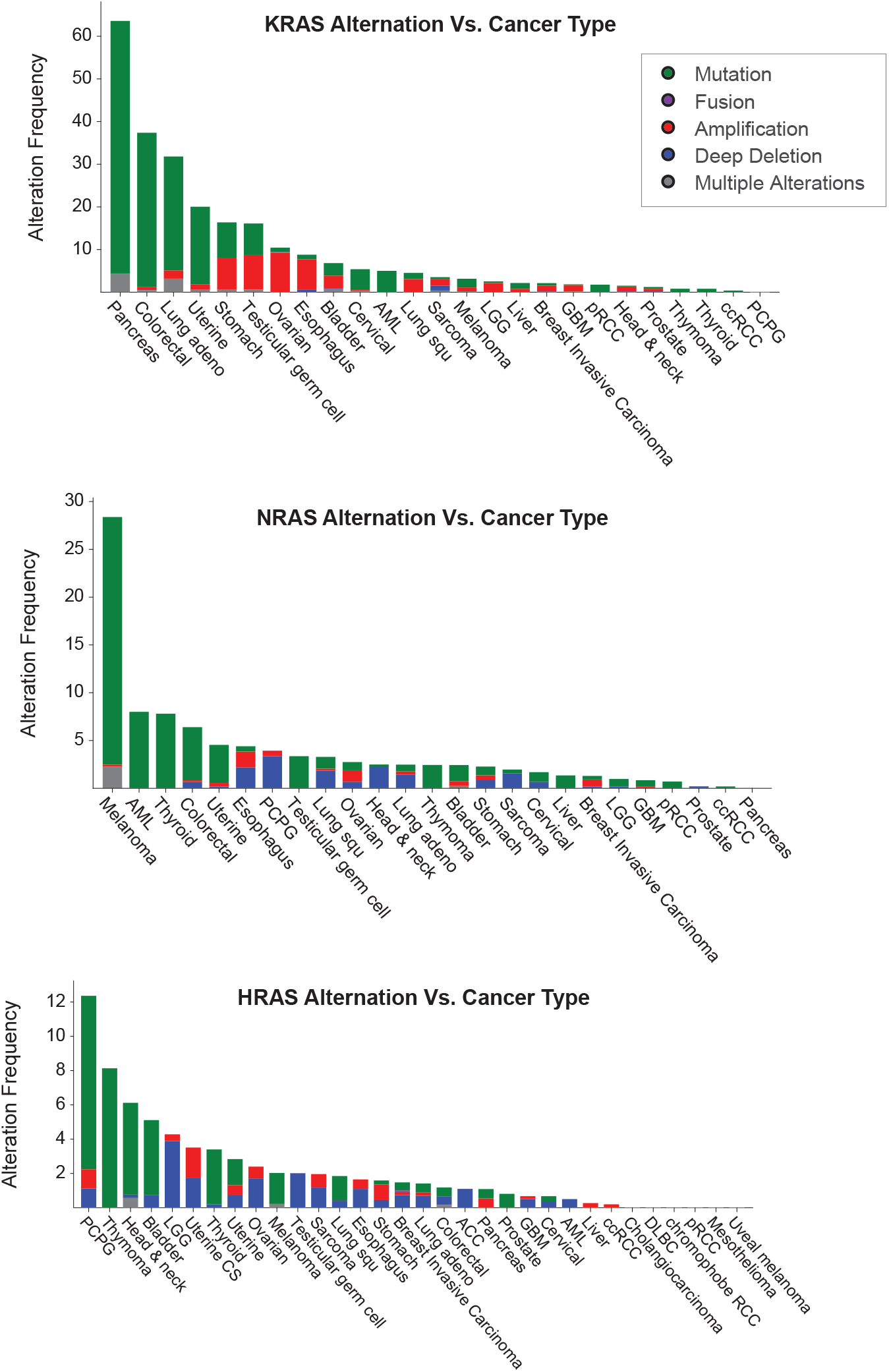
Alternation frequencies of KRAS (top), NRAS (middle), and HRAS (bottom) among different cancer types in the TCGA PanCancer Atlas cohorts. Frequencies of RAS protein alternation (i.e. mutation, fusion, amplification, deep deletion and multiple alternations) of different cancer types included in TCGA PanCancer Atlas cohorts were extracted via cBioPortal.

**Supplementary Figure 4.**
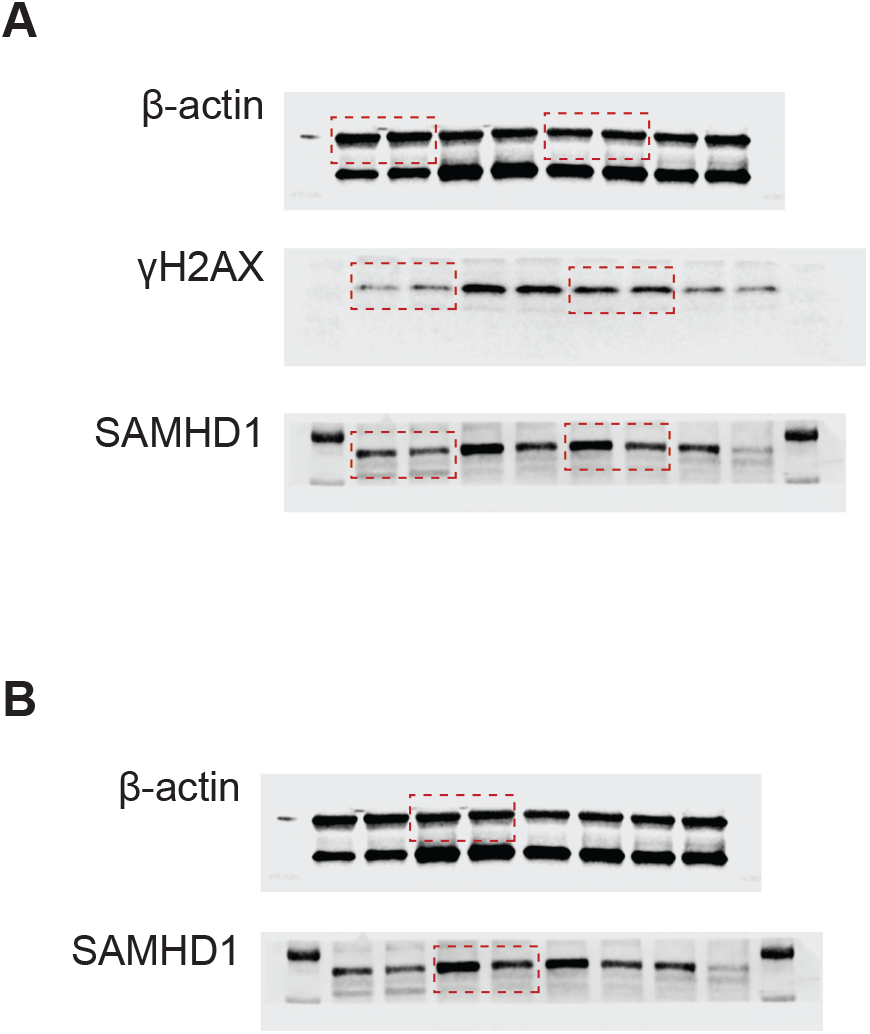
Uncropped Western blot images of Fig. 1D (A) and Suppl. Fig. 2A (B).

## Notes

### Competing Interest Statement

The authors have declared no competing interest.

